# A compact model for cold tolerance entrainment based on the ICE1-CBF-COR pathway in plants

**DOI:** 10.1101/2023.02.23.529825

**Authors:** Ting Huang, Yue Wu, Hengmin Lv, Yiting Shu, Linxuan Yu, Haoyu Yang, Xilin Hou, Xiong You

## Abstract

Most temperate plants can tolerate both chilling and freezing temperatures. Plants have developed self-defense mechanisms to perceive cold signal, trigger the cold tolerance for the cold response ICE1-CBF-COR pathway by packing a cascade of kinase phosphorylation/dephosphorylation events into a functional module. Linked to the central ICE1-CBF-COR model, two sub-models were adopted for the second messenSger Ca^2+^ -- a one-compartment model for Ca^2+^ transient and a two-compartment model for a train of Ca^2+^ spikes. Numerical simulation verified the expression patterns of the cold-induced genes as observed in *Arabidopsis thaliana, Brassica napa* and *Solanum lycopersicum*, including wild types and mutants. Moreover, the desensitization and resensitization of *CBF3* and *COR15A* were displayed as well as the dynamics in the gradually decreasing temperature response to cold stress. The duration of cold tolerance was predicted for approximate 10 days. Interpreting and predicting the dynamical behaviors of the cold signaling pathway are valuable and time-saving for understanding mechanisms of cold acclimation.

**Author summary:** In higher organisms, *CBF3* transcriptional level are output in a pulse and *COR15A* transcriptional level are maintained at a high homeostatic level to respond cold stress, which is determined by a complex set of regulatory mechanisms. Feedback loops from Ca^2+^ perception onto CaM, kinases, cold-acclimated genes in sequence, which have been experimentally reported, are thought to induce *COR15A* transcript accumulation under cold temperature exposure. The cold response pathway has been modeled based on a computational model. However, protein kinase cascades made the model relatively complicated and the cold-response gene *COR15A* was absent. Here, we develop a compact model with cold tolerance target gene *COR15A*. We show that this model could reproduce the temporal dynamics and characteristics of the core gene variables in different plants with wild type and mutant. Reduction of CaM proportion and augment of negative feedback control by ZAT12 in the model results in the desensitization of *CBF3* and *COR15A*. Less than 24-hr warm treatment loses the ability of resensitization of *CBF3* and *COR15A* after 14-day cold exposure. We further predict that the duration of cold tolerance is maintained at about 10 days in our model.

## Introduction

Low temperature stress is a negative factor that limits plant growth, spatial distribution and crop production [1]. In the long history of coping with environmental temperature changes, plants have developed a set of dedicate mechanisms to respond to cold stress and survive in low temperature conditions. This process is known as cold acclimation [2,3]. In the past decade the investigation of the molecular mechanism of cold acclimation has attracted increasing interest [4,5]. Recent advances have led to the understanding of plant cold-signaling perception and transduction [4,6–11].

Among several cold signaling pathways ever identified, the CBF centered pathway ICE1–CBF–COR has been acknowledged as the key regulatory element for cold stress response in plants [12,13]. When a plant is exposed to low temperatures, an increase in membrane fluidity and rearrangement of the cytoskeleton is caused, followed by the activation of calcium channels. A sharp influx of calcium triggers downstream responses [14,15], where three CBF/DREB1s (CBF1, -2, -3) are involved in the induction of the expression of cold related genes *CORs* [16,17]. The expression of *CBF/DREB1* (mainly *CBF3/DREB1A*) is in turn controlled by a MYC-type transcription factor ICE1 (Inducer of CBF Expression1) [18]. The expression of these *COR* genes, notably *COR15A*, enhances the cold tolerance through a variety of cellular regulatory mechanisms [19].

Apart from *Arabidopsis*, the *CBF* gene family and genome-wide characterization have been identified in many other chilling-sensitive plant species, such as non-heading Chinese cabbage [20], *Brassica rapa* [21], *Brassica napus* [22], tomato (*Solanum lycopersicum*) [23], cucumber [24], potato (*Solanum tuberosum*) [25], rice (*Oryza sativa*) [26] and wheat (*Triticum aestivum*) [27]. Thus, cold signaling pathway generally resides in different plants.

However, due to the complexity of the cold signal pathway where many chemical reactions and cellular processes are involved as well as the particularity of plants, experimental exploration is usually costly and time-consuming. As a result, mathematical models have employed to simulate the chemical kinetics of the plant cold signaling pathway. Very recently, based on experimental observations in *Arabidopsis*, a computational model of the ICE-CBF pathway was constructed to understand the molecular mechanism of the cold signaling in plants [28]. The model successfully recovered qualitative observations of plants responding to a single cold shock, both in wild type and mutants.

According to the above-mentioned study, many results obtained by research is desired to affirmative, but there are still some deficiencies. In the network structure, the inclusion of the cascade of kinase phosphorylation/dephosphorylation makes the model rather complicated, which consists of 14 ordinary differential equations with 77 parameters. From mathematical modeling point of view, the detailed description of kinetics of Ca^2+^-activated kinases, such as Ca^2+^/calmodulin-regulated receptor-like kinase (CLK1) and mitogen-activated protein kinases (MPK6, -4) is not necessary for modeling the core ICE1-CBF component. With respect to variable selecting, the important cold-related genes *CORs* were missing from the model, which prevented the model from verifying the experimental data on *COR* expressions and accurately measuring the cold tolerance of plants. Ultimately, part of experimental validations through model were not achieved such as the phenomenon of resensization.

Hence it is necessary to construct a regulatory model within core cold-responsive genes in plants. Using a combination of theoretical and numerical approaches, we showed that the expressions of main cold-related genes observed in different plants were verified by previous experimental data. The compact cold signaling pathway is characterized by the *CBF* centered network in a train of interlocking transcriptions and translations (Figure 1). By modifying kinase-regulated processes in a recent computational model, we simulated both different expression patterns between desensitization and resensitization of *CBF3* and *COR15A* level. Some predictions of this new model were more convincing than that in the previous one, such as *CBF3* and *COR15A* resensitization. Our simulation showed that the peak levels of *CBF3* mRNA were decreasing whereas of *COR15A* mRNA were increasing along with incremental warm/cold cycles. Plant, exposed to cold stress for 14 days after 2-warm treatment, needed 24-hr continuously heating to reproduce high expression of *CBF3* and *COR15A*, resulting in acquiring cold tolerance again. The duration of cold tolerance could be scaled by the continuous high expression time spans of *COR15A*, which The plant could withstand the cold stress for about 10 days.

**Fig 1.**
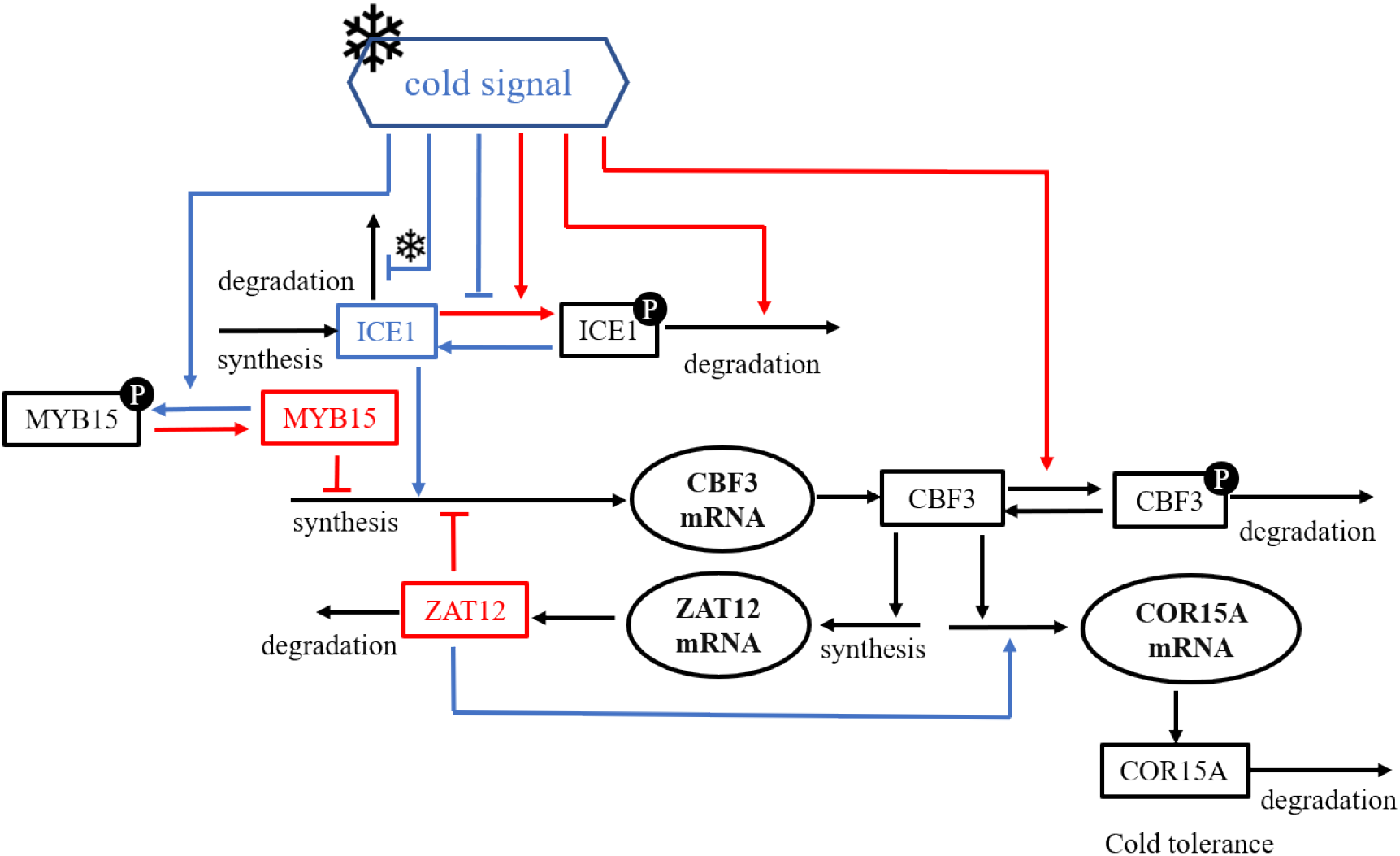
Scheme for the cold response pathway in plants. The low temperature (marked by a snowflake) contributes to a transient increase in Ca^2+^, in two forms of a single Ca^2+^ pulse or repetitive Ca^2+^ spikes. ICE1–CBF3–COR is represented as the core cold response pathway. The blue and red regulatory lines signify the positive and negative regulation of COR15A mRNA in the entire gene network, respectively. All the kinases (CRLK1, CRPK1, MPK6, MPK4, HOS1, 14–3– 3) in the model are grouped into a kinase module, and the regulation of downstream genes remains unchanged [28]. CaM activated by Ca^2+^ regulates downstream genes through the kinase module.

## Review of the cold response ICE1-CBF-COR pathway

In the past two decades, the gene regulation in plant cold signaling pathways has been extensively investigated by experiments [3,5,29,30]. ICE1-CBF-COR is the core variable in the cold response pathway.

### Cold signal perception

Ca^2+^ has been acknowledged as a key regulator with versatile functions in signaling pathways of many plants [31]. In plants, calcium signals are characterized by transient increase in cytosolic free Ca^2+^ ([Ca^2+^]_cyt_) under cold stress. The free Ca^2+^ is required to be sustained at a low level [32]. The activity of [Ca^2+^]_cyt_ results from the flux of Ca^2+^, the external medium and subcellular compartments where the concentration of Ca^2+^ is higher than that in the cytosol. Cold-induced [Ca^2+^]_cyt_ rising in plant cells sometimes occur as a pulse and sometimes present as repetitive oscillations or spikes, where the frequency (period), amplitude and shape (e.g. sinusoidal, square-wave) of the increasing curves depend on the essential character of the stimulus. Martin and Pittman showed that the resting [Ca^2+^]_cyt_ was stored 100 nM approximately after Ca^2+^ effusing [33].

Three main proteins, calmodulin (CaM)/CaM-like proteins, calcium-dependent protein kinases (CDPKs), and calcineurin B-like proteins [34–36], are identified to be Ca2+ sensors in plants. CaM, a Ca^2+^-modulated protein, has been proved to be a calcium-binding protein in all eukaryotes [33]. CaM has the function to regulate diverse downstream targets leading to a shift in physiology or developmental pattern under a variety of stimulus, including salinity, heat, cold, pathogens, hormone stresses, etc. [37– 39].

### Regulation of *COR* genes by CBF-dependent pathway

The greatest recorded genetic response pathway leading to gene induction under low temperature has been testified as the one mediated by C-repeat/dehydration-responsive element binding factors (CBFs) in different plants [40]. In the core cold pathway (ICE1-CBF-COR), the ICE1 acts as a positive upstream regulator of *CBF* leading to decreasing cold sensitivity. The expression of *COR* (cold-responsive) genes, controlled by CBFs, is vital for both freezing stress tolerance (below 0°C) and chilling stress (0–15°C) in plants [13]. Except for the basic component of the *ICE, CBF*, and *COR* genes, some other activators or repressors are involved in this pathway that directly or indirectly contribute to cold tolerance. Agarwal et al. [41] have demonstrated that the transcription factor MYB15 repressed the CBF3 mRNA level and reduced cold tolerance in *Arabidopsis*. The zinc-finger protein ZAT12 plays a crucial role in cold stress and acts as inhibitor of CBFs [42,43]. However, MYB15 has diverse functions on *CBF* transcript in different plants, such as a negative transcription factor in *Arabidopsis* and a positive regulator in tomato [44], where the experimental results showed that the exposure of tomato plants to cold temperature (4°C) increased the transcription levels of MYB15 and CBFs. In the model plant *Arabidopsis*, the phosphorylated MYB15 detaches from CBF3 promoters on account of reducing DNA-binding affinity [45].

The expression of *COR* genes is recognized to be necessary for plants to protect against cold stress. Many homologous *COR* genes have been identified in different plants, such as *COR6*.*6, COR47* in *Arabidopsis thaliana* [46], *COR25* in *Brassica napus* [47,48], and *WCOR15* in wheat [49]. The expression of the cold-regulated *COR15A* gene of *Arabidopsis thaliana* results in a significant increasing survival in freezing and chilling stress [50]. Therefore, *COR15A* gene was chosen as a representative gene against cold tolerance.

## Results

### Dynamic verification of cold-related genes under Ca^2+^ one-compartment model

CBFs, notably CBF3, is demonstrated to regulate the expression of *COR* genes, which are activated to cope with cold stress [51]. The expression level of *CBF3* and *COR15A* mRNA were core measured variables respond to low temperature in the model. The model was no longer concerned on the specific kinases regulating the core genes in the downstream of cold response pathway, but only focused on the positive and negative regulation of the downstream genes by Ca^2+^ influx, mediated by CaM, through the kinase module. The model complexity was greatly reduced (numbers of variables and parameters) while preserving the better performance.

The time courses of the cold response variables were elucidated (Figure 2) after the temperature drops to 4°C *a*t *t* = 2 h, in order of the proportion of CaM (A), the concentration of ICE1 (B), MYB15 (C), *CBF3* mRNA (D), *ZAT12* mRNA (E) and *COR15A* mRNA (F). Experimental studies focused on the quantification of *CBF3* mRNA, *ZAT12* mRNA and *COR15A* mRNA [52–54]. Comparison of model prediction results and experimental data was shown in the second row (Figures 2D, 2E, 2F), where black solid curves, red, blue, and green dots represented the relative abundance of simulation, experimental data for *CBF3* mRNA, *ZAT12* mRNA and *COR15A* mRNA. We continued to integrate the time evolution of Ca^2+^, ICE1, MYB15, *CBF3* mRNA, and *ZAT12* mRNA (Figure S1) to obtain systematically understanding the dynamics of the cold response pathway. The different temporal processes in the same diagram again showed the order of gene activation in the cold response pathway.

**Fig 2.**
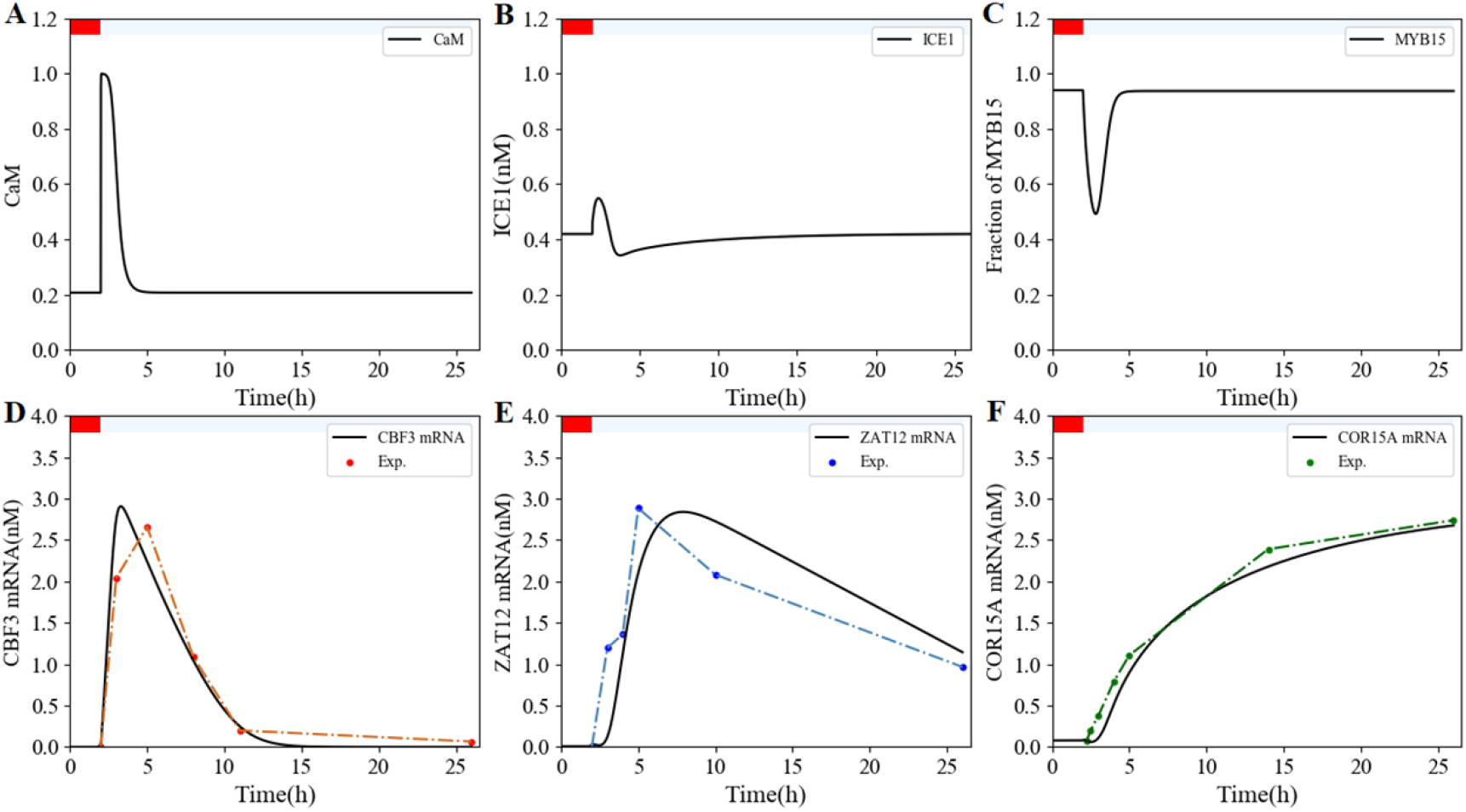
Dynamics of the core cold response pathway under the cold stimulation. (A). The cold stress causes Ca^2+^ to increase instantaneously at ZT2. This single Ca^2+^ pulse gives rise to transient changes in gene expression (ICE1 (B), MYB15 (C), *CBF3* mRNA (D), *ZAT12* mRNA (E) and *COR15A* mRNA (F)) throughout the network. Experimental data (red dots for *CBF3* mRNA (D), blue dots for *ZAT12* mRNA (E), green dots for *COR15A* mRNA (F)) is derived from Figure 1 in Zarka et al. [65], from Figure 7A in Fowler and Thomashow[53] and from Figure 3(a) in Kang et al.[54]. Black solid curves denote the simulations. The results were obtained by numerical integration of kinetic equations (4) – (13), by the classical fourth-order Runge-Kutta method. The kinetics of Ca^2+^ and activated CaM are the same as that in computational model (see Sections 1 in Supplementary Material). Parameters are showed in Supplementary Table 2. The red bars represent warm temperature treatment, and the aliceblue bars indicate low temperature treatment.

The concentrations of *CBF3* mRNA (Figure 2D), and *ZAT12* mRNA (Figure 2E) showed a rapid growth followed by a descent and the transcription level of *COR15A* demonstrated a sustained rising in 24-hr (Figure 2F) after low temperature stress. The simulated relative abundance was well fitted with the experimental data. The transient Ca^2+^ influx led to the moment changed variables in the cold response pathway. Ca^2+^ concentration reaching a peak within a few minutes returned to a steady state that prevailed prior to the cold stress (Figure S1), while downstream genes had a few hours or some 10 hours to respond (Figure 2E, Figure S10D). After Ca^2+^ pulse, MYB15 reached its trough, followed by the peaks in *CBF3* mRNA and *ZAT12* mRNA. *CBF3* mRNA returned to steady state earlier than *ZAT12* mRNA, due to the relatively slow degradation of *CBF3* mRNA under low temperature.

The same result as the previous computational model [28] was that the concentration of ICE1 and MYB15 raised and dropped first respectively and then returned to equilibrium (Figure 2B, C). That is because the transient increase of Ca^2+^ concentration under cold stress causes phosphorylation of MYB15, resulting in a decrease in the amount of MYB15. When the amount of MYB15 reached its trough, the inhibition of CBF3 was weakened and the expression of *CBF3* mRNA increased. The concentration of ICE1 increased in time after a drop in temperature owning to the constant synthesis of the ICE1 protein at a rate *k*_*s1*_ *i*n eq. (5), and then decreased below the initial value. As time grows, the concentration of ICE1 eventually tended to the steady state equal to the initial value.

In order to verify the versatility, the model was applied to other temperate-sensitive vegetable plants, including tomato (Figure 3B, 3E) and *Brassica rapa* (Figure 3C, 3F). Individual parameters were appropriately modified in simulation so that the simulation results fitted well with the experimental observations. Comparing with a continuous increasing of *COR* genes in *Arabidopsis* and *Brassica*, the expression of cold-related gene *SlCOR413* went up followed by a small decline in 24-hr. *Arabidopsis* and *Brassica* are members of the cruciferous family, while tomato belongs to the Solanaceae family.

**Fig 3.**
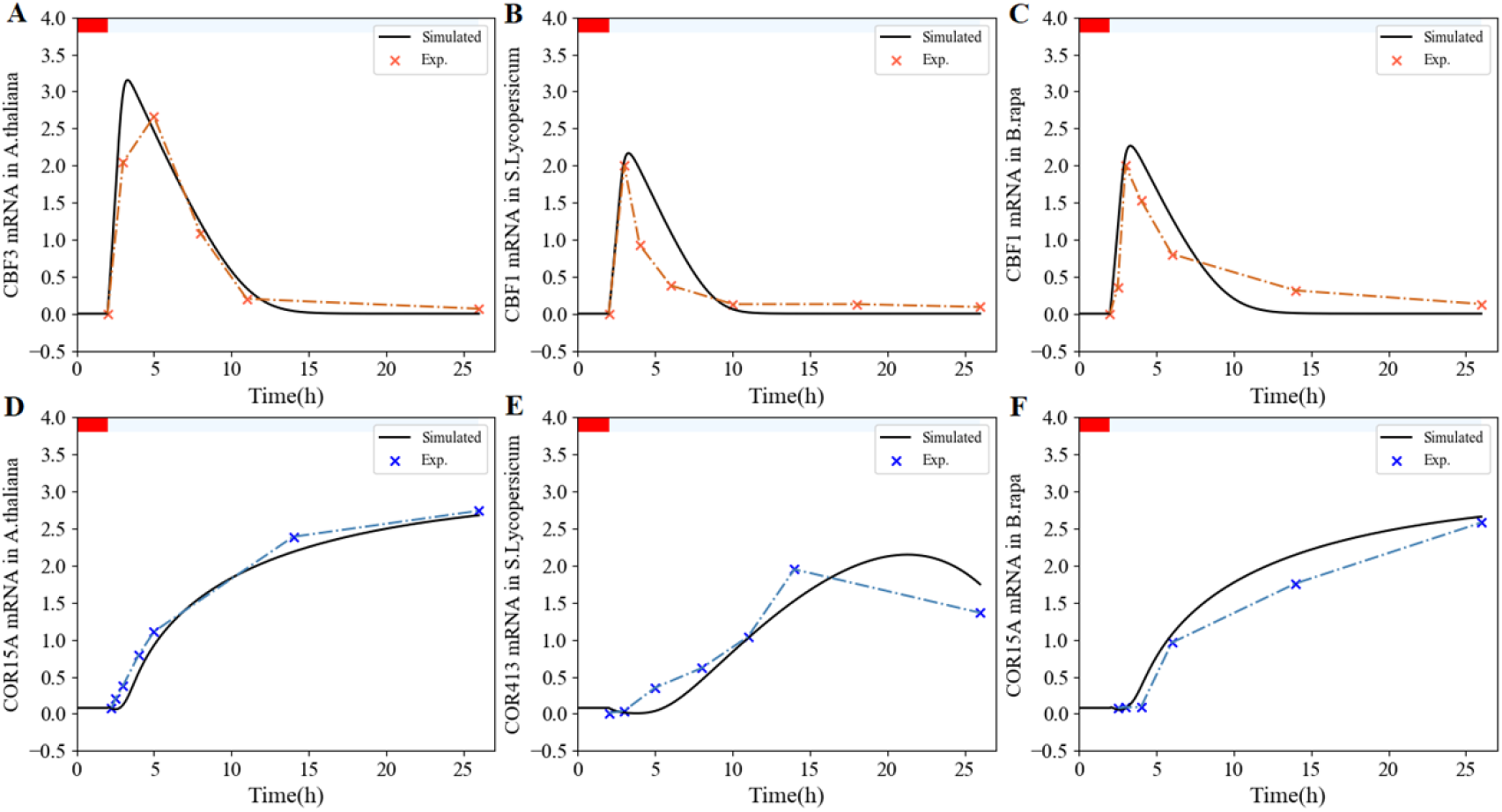
Dynamics of *CBFs* and *CORs* in the compact cold response pathway in different plants induced by the one-compartment Ca^2+^ model compared to the corresponding experimental data. Diurnal variations in gene expression of *CBFs* and *CORs* mRNA occur in *Arabidopsis thaliana* (A and D), *Solanum lycopersicum* (B and E), and *Brassica rapa* (C and F) are simulated, comparing to the experimental observation in Zarka et al. [65], Wang et al. [90] and Wang et al. [20], respectively. Parameter values are listed in Supplementary Table 2 except for *v*_*s1*_ *= 30 n*M/h, *v*_*s2*_ *= 8 n*M/h, *v*_*s3*_ *= 1*.*4 n*M/h, *K*_*I2*_ *= 0*.*7 n*M, *K*_*I5*_ *= 1*.*8 n*M, *K*_*I6*_ *= 4 n*M, *v*_*d7*_ *= 1*.*6 n*M/h, *K*_*d3*_ *= 0*.*4 n*M, *K*_*d6*_ *= 1*.*5 n*M in *Solanum lycopersicum* and *v*_*s1*_ *= 25 n*M/h, *v*_*s2*_ *= 8 n*M/h, *v*_*d7*_ *= 0*.*4 n*M/h in *Brassica rapa*. The red bars represent warm temperature treatment, and the aliceblue bars indicate low temperature treatment.

The time evolutions of the core variables in the cold response pathway provided a qualitative explanation for the regulation mechanism. The simulated results showed us a brief view of the process from the one-compartment Ca^2+^ model to the synthesis of cold response genes.

### Dynamic analysis and prediction of cold-related genes under Ca^2+^ two-compartment model

The concentration of Ca^2+^ in cytoplasm ([Ca^2+^]_cyt_) is transiently elevated to avoid toxicity, and transmitted to distant cells or organs in the form of a single spike, oscillations or waves [55,56]. The duration, periodicity and amplitude of [Ca^2+^]_cyt_ oscillation were measured in *Arabidopsis* guard cells with an amplitude of about 125 nM and a period of about 150 s [57,58]. [Ca^2+^]_cyt_ maintains low levels in resting or unstimulated cells, which is suitable for cytoplasmic metabolism. Localized [Ca^2+^]_cyt_ oscillations were generated through enzymes’ interplay with Ca^2+^ channels in cold stress [59]. The Ca^2+^ influx from the apoplast, proportional to parameter *β (*red curve in Figures S2A, B), still adopted the same dynamic equation (see eq. (2) in supplementary) in the previous computational model [28,60,61].

The exponential decrease after the transient increase of *β s*till led to a series of Ca^2+^ transient oscillations (blue curves in Figures S2A, B). The expression of *CBF3* mRNA was simulated by the fourth-order Runge-kutta method [62] corresponding to the train of transient spikes above (Figures S2C, D). The simulation results indicated that the duration of the Ca^2+^ oscillation affected the accumulation of *CBF3* mRNA and the duration of the Ca^2+^ spike sequence relied on the decay rate of *β*.

Interestingly, the parameters *β*_*f*_ *a*nd *a c*ould affect the spike behaviors of Ca^2+^. The ratio of *a t*o *β*_*f*_ approximately less than 25 resulted in multi-spike oscillation of Ca^2+^, while in the form of single spike as the ratio more than 25 (Figure S3A). The special oscillation pattern (Figure S3B) and single spike mode (Figure S3C) of Ca^2+^ were obtained by taking any point in the multimodal and unimodal region, respectively.

### Effects of mutation and overexpression in the cold stress pathway

The cold-activated model retained the predictive function of influence mutations and overexpression. The transcriptions of *CBF3* were analyzed as a marker, with ICE1, MYB15 mutant and ZAT12 overexpressed. In order to represent these mutations, the synthesis rate (*k*_*s1*_) of ICE1 was reduced to 20% of the original value (Table S2). The dephosphorylation rate (*v*_*1*_) of phosphorylated MYB15 was reduced to 10%. The synthesis rate (*v*_*s2*_) of ZAT12 was increased by 5-fold, indicating its overexpression. The results were presented in Figure 4 (black and blue solid lines denote WT and mutants/overexpression, respectively; black and blue dots represent experimental data, respectively).

**Fig 4.**
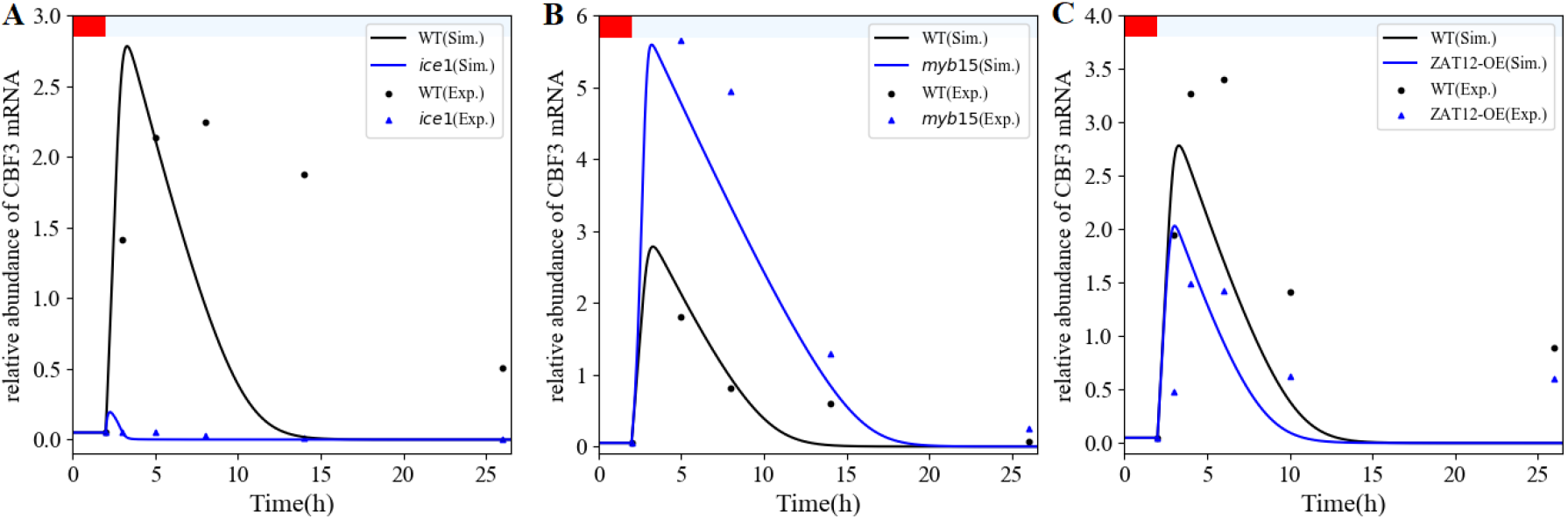
Comparing the effect of gene mutation and overexpression between experiment and simulation. The black and blue solid lines represent the simulated distribution of *CBF3* mRNA expression in wild-type and mutant or overexpression ((A) the *ice1* mutant, (B) the *myb15* mutant, (C) the overexpression of ZAT12 (ZAT12-OE)), the black and red solid dots indicated as the experimental data in each panel. In (A), the synthesis rate (*k*_*s1*_) of ICE1 in the *ice1* mutant is 0.036, compared to 0.18 in WT (Table S2). Experimental data are extracted from Figure 1C in Chinnusamy et al.[52], by means of the software ImageJ (https://imagej.nih.gov/ij/). In (B), the dephosphorylation rate (*v*_*1*_) of phosphorylated MYB15 protein in the *myb15* mutant is 0.4, 4 matched in WT. Experimental observations are redrawn from Figure 7C in Agarwal et al.[41]. In (C), the maximum synthesis rate (*v*_*s2*_) of ZAT12 mRNA is 47.5 versus 9.5 in WT. Experimental data are obtained from Figure 10A in Vogel et al.[43]. Other parameters are the same as that in Figure listed in Table S2. The red bars represent warm temperature treatment, and the aliceblue bars indicate low temperature treatment.

The model allowed us to simulate the fact that the *ice1* mutation inhibits *CBF3* transcription (Figure 4A), which was remarkably increased by ICE1 overexpression (ICE1-OE; Figure S4) in accord with experimental observations [52]. MYB15, a repressor of *CBF3* transcription, remained inactivated by phosphorylation in low temperature. Transcriptional repression of *CBF* genes and the cold tolerance were mediated by direct phosphorylation of MYB15 [41,45]. Therefore, MYB15 mutation directly led to an increase in CBF expression (Figure 4B; Figure S5).

ZAT12 was confirmed to downregulate the transcription of the *CBF* genes, while CBF3 protein upregulating the expression of *ZAT12* mRNA, thus forming a negative feedback loop [1,63]. About 2 h after cold stimulation, the level of *CBF3* mRNA under ZAT12 overexpression coincided with the corresponding in wild type (Figure 4C), because the amount of *ZAT12* was low at room temperature and it took a phase of time for *ZAT12* transcription by CBF3 protein in cold stress.

### Cold-sensing mechanism of *CBF3* and *COR15A* transcript (desensitization and resensitization)

Plants experience a variety of external mechanical stimuli and adapt to their growth and development by triggering a series of signal events. In the past two decades, the research of plant mechanical sensing has attracted more attention, but plants response to repeated mechanical stimuli is not described in a detailed model.

Quantifying plant mechanical stimulations and comparing the effects of different degrees or amounts of these stimulations were general approaches in the past. For example, decay of Ca^2+^ peaks is observed and desensitization starts in several minutes after repeated increasing and decreasing temperature [64]. The same phenomenon can be seen at the molecular level as the gene expression was rapidly modified by exposing plant to low temperature. It is reported that the cold-sensing mechanism, represented by the accumulation of CBF expression is desensitized within a few hours [65].

The resensitization mechanism has been evaluated different from the one leading to desensitization [66]. Knight et al. [67] investigated that wind-induced stimulations of Nicotiana seedlings caused an increase in the peak of [Ca^2+^]_cyt_. However, if the stimulation was repeated every 5 seconds, the amplitude of the peak would decrease. Complete desensitization was achieved after approximately 6 or 7 repetitive stimuli. The resensitization reaction would occur after 60 seconds with stimulation decreasing.

The previous studies allowed us to make simulated prediction comparing with the experimental results. For the sake of validation and prediction of system, a train of Ca^2+^ pulses were occurred in the warm/cold cycle as repetitive stimuli. The time courses of *CBF3* mRNA and *COR15A* mRNA were then quantified under warm/cold period of 1.5 h/1.5 h (obtained experimentally by Zarka et al. (2003), Figure 5A, 5D), 3 h/3 h (Figure 5B, 5E), 6 h/6 h (Figure 5C, 5F). The degradation rate of *CBF3* mRNA in warm conditions was assumed to be 10-fold that in cold conditions. Furthermore, in order to take the kinetic of resensitization into account, plants that have been cold-acclimated at 4°C for 14 days were returned to warm temperatures for different time spans, followed by warm/cold cycle presented in Figure 6.

**Fig 5.**
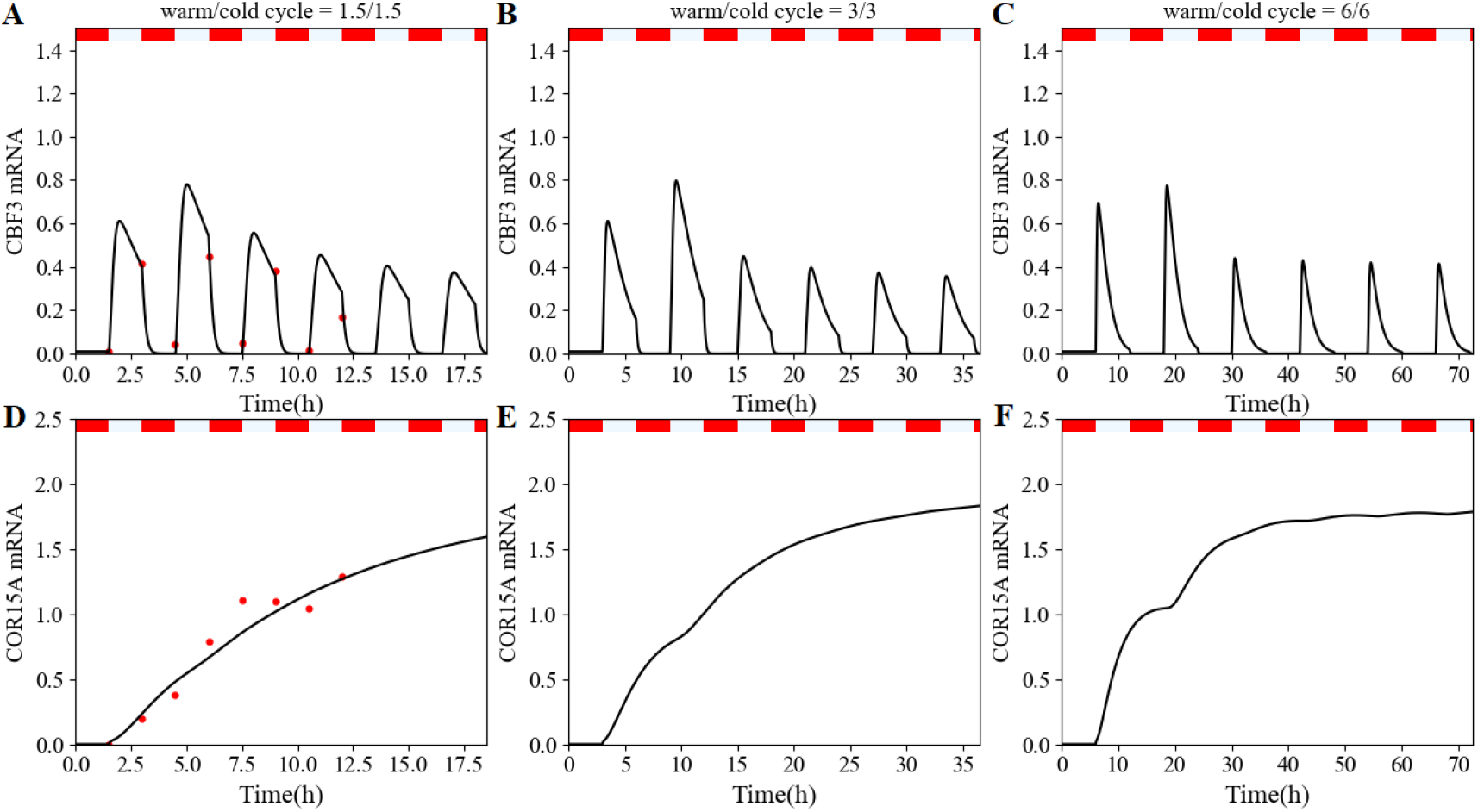
Dynamics of plant cell response to repeated cold stress stimulations. The expression distribution of *CBF3* mRNA and *COR15A* mRNA (black curves) under the uniform Ca^2+^ pulse (blue curves) is shown, allocated time intervals of 3 h (A, D), 6 h (B, E) and 12 h (C, F). The durations of Ca^2+^ pulses are identical in each panel. We tested 3 cycles characterized by a period of 3 h (90 min cold followed by 90 min warm, A), 6 h (B), and 12 h (C), respectively, under the assumption that each repeated cold stress produces the same Ca^2+^ pulse. Experimental observations (red solid dots) are extracted from Figure 2A in Zarka et al.[65]. The red and Alice blue bars over each panel signify the warm/cold (20°C/4°C) cycles, respectively. Numerical integration of eqs. (1) and (4) – (13) solved by the fourth-order Runge-Kutta method in the software python (https://www.python.org/), with different activation rate of calmodulin by Ca^2+^ (*K*_*Ca*_ *= 0*.5 vs. 0.14) and degradation rates of *ZAT12* mRNA (*v*_*d6*_ *= 0*.001 vs. 0.1). The red bars represent warm temperature treatment, and the aliceblue bars indicate low temperature treatment.

**Fig 6.**
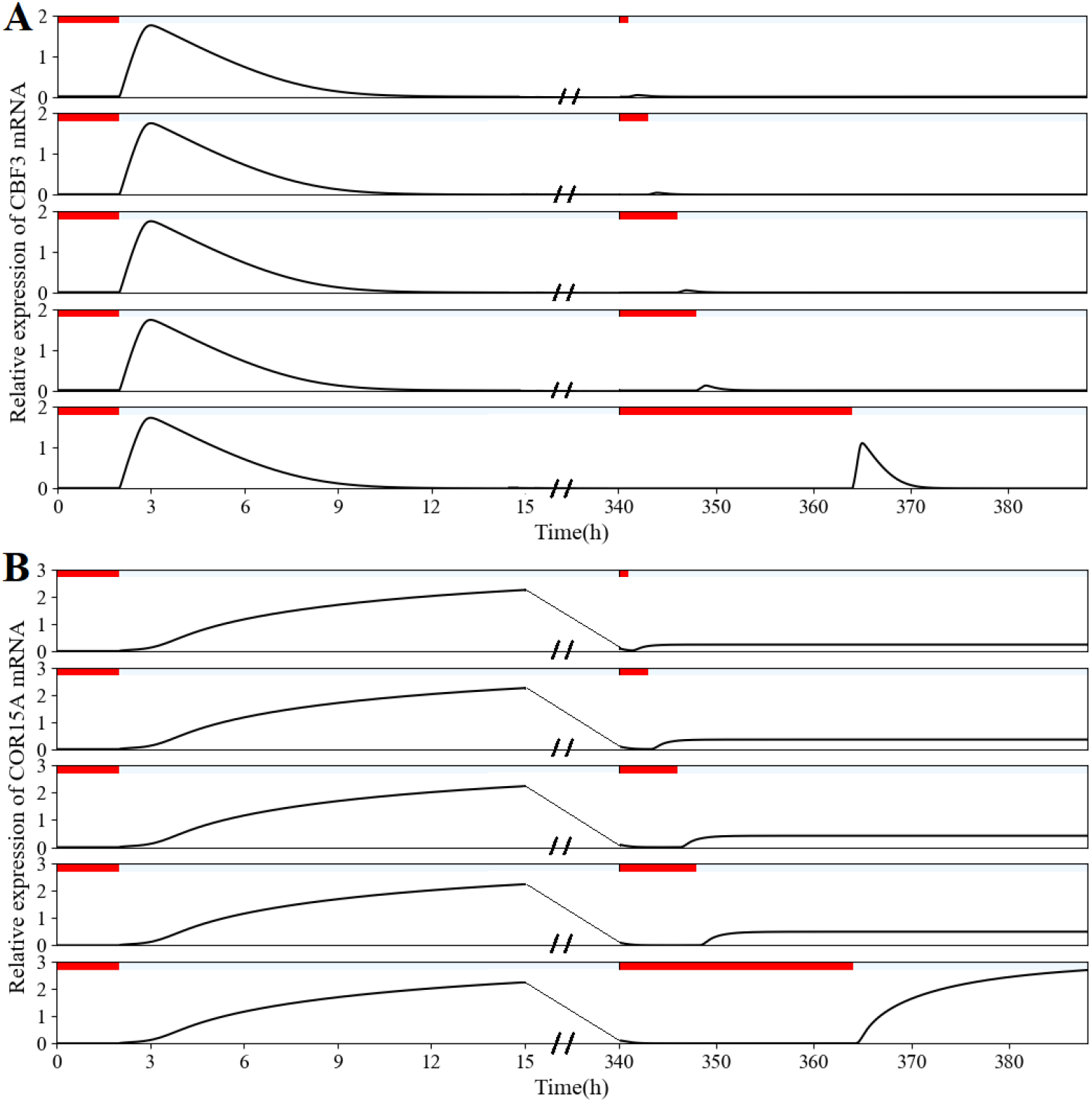
Dynamics of resensitization of the regulatory circuit controlling accumulation of *CBF3* and *COR15A* transcripts. An acclimated plant can response to cold stress after up to 24 h exposure in warm temperature (20°C). Experimental evidence can be found in the literature [65]. But *CBF3* (A) and *COR15A* (B) transcript levels depend on the time span of the warm treatment. In the experiment, plants were exposed to 2-hr warm treatment, followed by cold acclimated at 4°C for 14 days. After cold acclimation, plants were returned to warm temperatures at specified length of times (1 h, 3 h, 6 h, 8 h and 24 h), and then subjected to a cold shock at 4°C. The visualization of 14 d cold-acclimated was reduced (denoted as two slashes) as *CBF3* transcript levels maintained at a low equilibrium state and *COR15A* transcript increased to high steady-state level followed by decreasing to low equilibrium. The red and aliceblue bars represent warm and cold temperature treatment, respectively. Parameter values are listed in Table S2 except for *K*_*Ca*_ *= 0*.*31 (*panel A), *K*_*Ca*_ *= 0*.*18 (*panel B).

Numerical simulations show that *CBF3* mRNA levels augment with cold stimulation and reduce with plant kept in warm condition and the increasing degradation rate. The variations of *CBF3* mRNA levels are discrepant in different warm/cold cycles. The concentration of *CBF3* mRNA appears to be desensitized during under different warm/cold cycles, i.e., the corresponding amplitudes gradually diminished with repetitive cold stimulations. But the desensitization phenomenon was not visible obviously over time, especially in the 6-hr warm /6-hr cold cycle (Figure 5C). This result manifested that the desensitization phenomenon was related to the frequency of stimulation in a relatively constant time. In addition, the phases of *CBF3* mRNA under disparate warm/cold cycles were consistent with the one of Ca^2+^ pulse with the extension of the stimulation cycle. The repeated cold stimulation of the same intensity gradually increased the transcription level of *COR15A* to a steady state (Figure 5D, E, F), while the longer warm/cold cycle led to deep fluctuation of *COR15* mRNA. Therefore, the model proved that cold acclimation contributes to the improvement of cold tolerance of plants. Thus, mechanosensitive control is an adaptive trait beneficial to plant growth and development [68].

The model could also realize the resensitization of plants. In experiments, plants were acclimated at 4°C for 14 days and then were transferred to warm temperatures (20°C) for 1 h, 3 h, 6 h, 8 h, 24h and then subjected to a cold shock abruptly. There were few observable increases in *CBF3* and *COR15A* transcript levels after returning to 1 h, 3 h, 6 h, 8 h warm temperatures in simulated results (Figure 6). However, if cold-acclimated plants were allowed to expose to warm temperatures for 24 h and then transferred to 4°C, *CBF3* and *COR15A* transcript levels obviously increased, qualitatively consistent with the experimental data (Fig. 2B in [65]). The experimental data and numerical simulated results indicated that prolonged exposure to low temperatures made cold-sensing mechanism of plants become desensitized but become resensitized to 4°C after 24-hr at warm temperatures treatment. The resensitization process required approximately 8 to 24 hours of warm temperature treatment, as cold-acclimated plants that had been returned to warm temperature for 3, 6, and 8 h were unable to induce normal expression of *CBF3* transcripts.

Except for the desensitization caused by reduplicative warm/cold shocks, the dynamics of transcription levels of *CBF3* and *COR15A* were investigated under gradually decreasing temperature as well. The experimental finding and the numerical simulation both supported the conclusion that *CBF3* and *COR15A* transcript levels were slightly and strongly induced when the plant was exposed to a gradual decrease in temperature from 20°C to 10°C and from 10°C to 4°C, respectively. Consistent with the previous results (Figure 2, Figure S1), prolonged exposure to constant low temperature (10°C or 4°C) resulted transcriptional declines in *CBF3* and *COR15A*, i.e., transcription levels of cold-responsive genes were powerfully induced upon a new round of gradual cooling (Figure 7). Because diverse temperatures affected pump sensitization and desensitization interactions after plant cold shock, which altered the magnitude of the transient Ca^2+^ response and the value of the Ca^2+^ level [69]. Therefore, we distinguished the maximal calcium ion current that can be achieved under low temperature shock in the Ca^2+^ model, namely the parameter *I*_*INmax*_ was set to 20 and 30 cooled at 10°C and 4°C, respectively.

**Fig 7.**
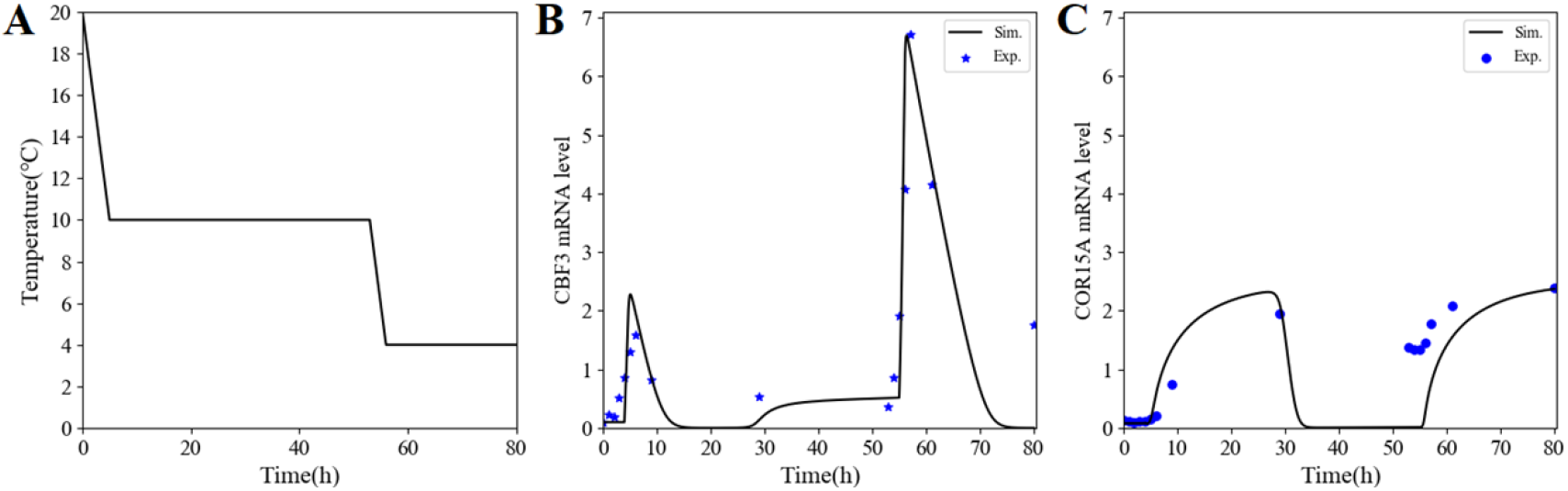
The dynamic of *CBF3* and *COR15A* transcript level under gradual temperature decrease condition. *CBF3* and *COR15A* transcripts in plants were accumulated when the temperature dropped from 20°C to 4°C. The color bar indicates the temperature decline at 2°C per hour. From ZT5, the plant was exposed to constant 10°C for 48 hours, followed by cooling to 4°C at the same decreasing rate, and then held at 4°C. The compact model was used to simulate the dynamic behavior in this particularly cooling mode in order to attain a better quality of experimental data recovery. Parameter values used for simulation are shown in Table S2 except for *K*_*Ca*_ *= 0*.*18*. Additionally, in order to distinguish the Ca^2+^ maximal input between cold shock at 10°C and 4°C, we set *I*_*INmax*_ *= 20 a*nd *I*_*INmax*_ = 30 cooled at 10°C and 4°C, respectively.

### Cold tolerance scaled by specific parameters

Model parameters have important roles on the steady-state durations of *COR15A* mRNA level (Figure 8). Increasing several rates, i.e., *v*_*2*_, *k*_*1*_, *K*_*1P*_, *K*_*2*_, *k*_*s1*_, *k*_*s2*_, *K*_*a1*_, *K*_*a3*_, *K*_*I2*_, *K*_*I3*_, *K*_*d2*_, *K*_*d3*_, *K*_*d4*_, *K*_*d5*_, *K*_*d6*_, led to high transcriptional level of *COR15A*, meanwhile maintaining high equilibrium could be achieved by decreasing other rates, i.e., *K*_*Ca*_, *v*_*1*_, *k*_*2*_, *K*_*1*_, *K*_*2P*_, *v*_*d2*_, *v*_*d3c*_, *v*_*d4*_, *v*_*d5*_, *v*_*d6*_, *v*_*d7*_, *K*_*I4*_, *K*_*d7*_, resulting in a longer duration of cold tolerance. Whereas six specific parameters, i.e., *v*_*d8*_, *v*_*d9*_, *K*_*I5*_, *K*_*I6*_, *K*_*d8*_, *K*_*d9*_, determined the duration of cold tolerance (∼10 days) through the model. These parameters involved in maximum rates of enzymatic degradation, threshold constants for activation, Michaelis constants for degradation of COR15A mRNA and protein.

**Fig 8.**
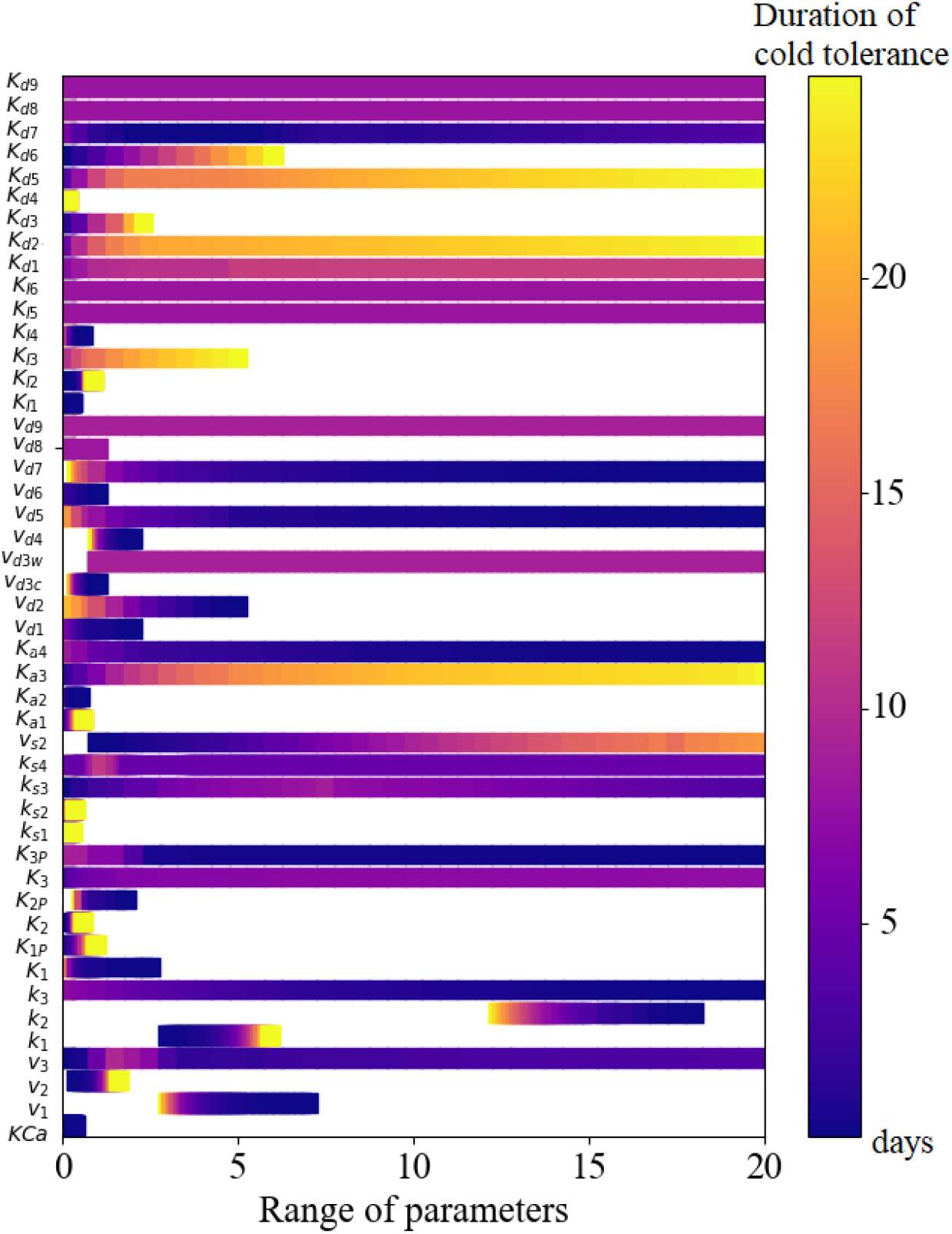
Cold tolerance was measured by the durations of *COR15A* maintained at a high steady-state level. We depicted the duration of cold tolerance through the continuous days of the steady-state of *COR15A* transcriptional level based on the variations of each parameter. The gradient from blue to yellow denote the sustained days of cold tolerance in plant.

### Presumptive oscillation of *CBF3* and *COR15A* induced by cold shock

The negative feedback loop is intensively deemed as primary cause for oscillation at the cellular level [70,71]. Interestingly, cold-induced gene oscillation has been predicted in the computational model published in 2020, due to the negative feedback of CBF-ZAT12 [28], but only for *CBF3*. In the dynamic study of plant cold response pathway based on the compact model, *CBF3* oscillation was recovered, resulting from the negative feedback exerted by ZAT12 under room temperature (20°C) as well (Figure S6). And the negative feedback loop located at the end of the whole network (Figure 1). The time series of mRNA (Figure S6A) and protein (Figure S6B) indicated sustained periodic oscillations, quantized by a closed limit cycle (Figure S6C). The duration period under the basic parameters and parameters in Figure S6 are approximately 22 h and 10 h, respectively. The detailed characteristics of the oscillation refer to the Figure S7 and Figure S8. In addition, consistent with the computational model, the numerical simulation reveals that ICE1, an upstream gene in cold response pathway, remained at steady-state both at constant room temperature (20°C) and from cold (4°C) to warm (20°C) temperature (Figure S6D).

Then, of course, there are the questions of whether gene oscillations occur if negative feedback emerged in the modelling, and whether oscillations caused at room temperature can also occur at low temperature? It is unexpectedly predicted that the transcription oscillation of *CBF3* and *COR15A* would occur after at least one day or two days under particular parameters upon chilling exposure (4°C) (Figure 9A), i.e., the negative feedback loop is not a unique factor for the oscillation of *CBF3* and *COR15A* level. We explored the key features (period, the first peak time, amplitude) that governed the periodic rhythmicity of cold-induced genes via series of random parameterizations which included the parameter *K*_*Ca*_ *c*losely related to the proportion of CaM activated by Ca^2+^, synthesis constant rate *v*_*s1*_ *a*nd degradation rate *v*_*d4*_. Our modelling analysis demonstrated that continuous oscillation of cold-related genes could likewise be induced at low temperature within a specified range of parameters. The black dots in Figure 9B denoted the periods of *CBF3* and *COR15A* mRNA levels, which increased with the increasing values of *K*_*Ca*_. Meanwhile, phase advance of *CBF3* transcript (blue dots) and phase delay (green dots) of *COR15A* transcript were clearly observed with the incremental variation of *K*_*Ca*_. Figure 9C and 9D summarized the main results concerning amplitudes of the above cold-responsive genes in changing parameter *K*_*Ca*_.

**Fig 9.**
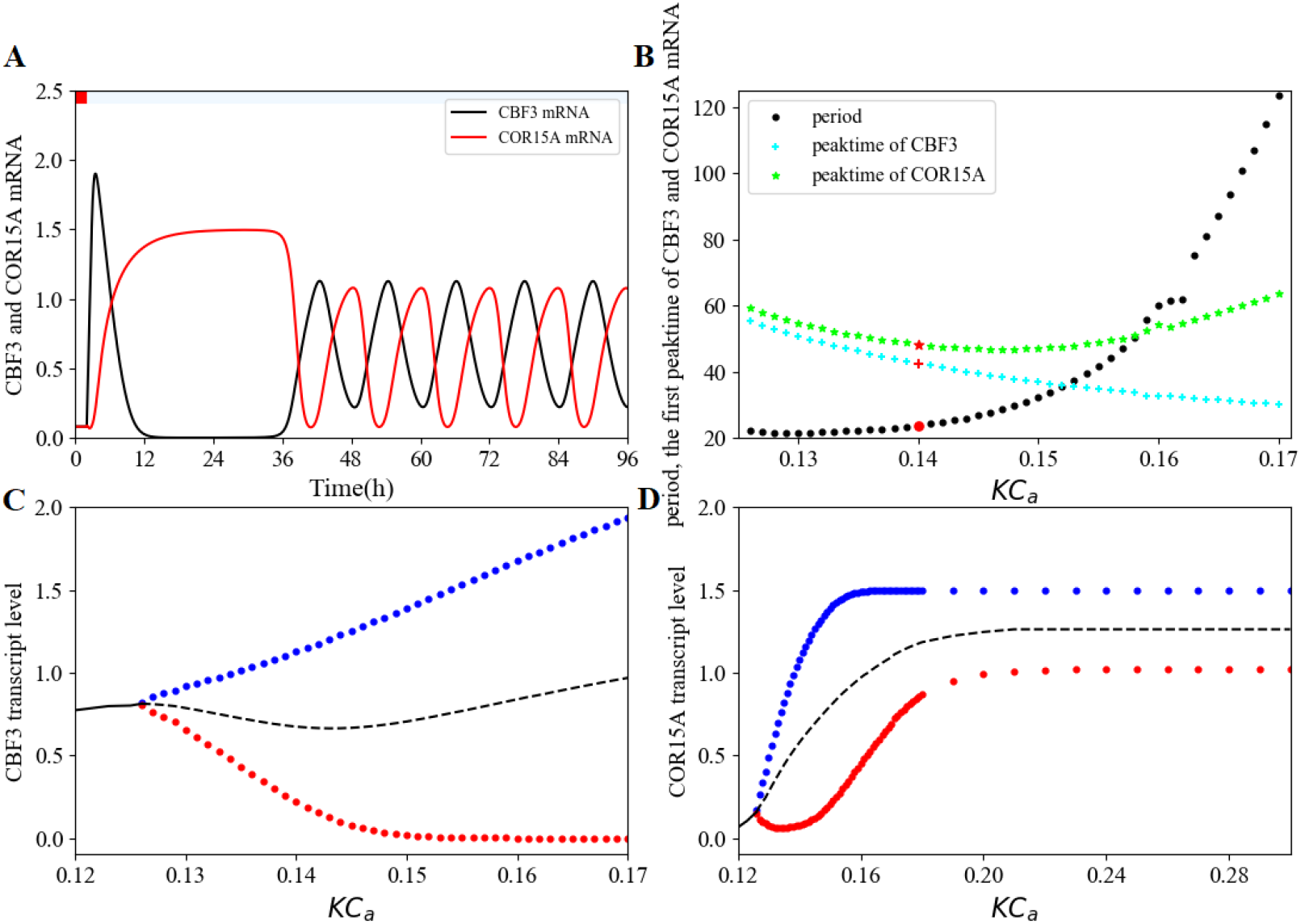
Temperature-induced oscillation of *CBF3* and *COR15A* transcript with specific parameters. Except for CBF3-ZAT12 negative feedback loop, other avenues leading to cold-responsive gene oscillation in cellular level were also investigated. A, based on the basic parameters, continuous oscillations was generated by substituting *v*_*s1*_ *a*nd *v*_*d4*_ *w*ith 20 (34 originally) nM/h and 0.9 (1.2 originally) nM/h, respectively. Moreover, the degradation rate (*v*_*d8*_) of COR15A mRNA was 1.5 and 1.25 in cold and warm phases, respectively. As an important parameter linking Ca^2+^ and calmodulin, *K*_*Ca*_ *w*as focused to discuss its effect on the period (black dots in panel B), first peak time (blue and green dots in panel B), and amplitude of *CBF3* (C) and *COR15A* (D) transcript level.

Furthermore, under the same set of parameters as Figure 9A, we compared the time evolution of *CBF3* and *COR15A* mRNA in the single-spike (Figure 9A) and the multi-spike Ca^2+^ model (Figure S10). The time series modeled in Figure 9A and Figure S10 provided an interesting representation of the variations on oscillation of the different Ca^2+^ mechanisms involved in the cooling response. Thus, the time courses under one set of parameters respond to periodic rhythm associated with the waves of Ca^2+^. The controlled simulations signified that negative feedback, parameters variation and temperature change were all likely to induce system oscillations.

## Discussions

Understanding how plants acquire cold acclimation is valuable for the improvement of the cold tolerance in plants and for anticipating the effects of climate variation on crop yield. Our investigation aims to establish a compact model, widely applicable to cold tolerance in a series of plants. The validity of the model is identified by comparing with computational model and experimental data [72,73]. In addition, the effectiveness also could be validated in *Arabidopsis* and horticultural crop (tomato and Chinese cabbage; see Figure 3). The model underlines transduction process of the initial cold signal through successive steps. The model simplifies the theoretical framework of the computational model and retains the core ICE1-CBF-COR regulatory pathway. The molecular characteristics and mechanisms of disparate plants response to low temperature are different, but the peaks of cold-activated *CBFs* gene range from 1h to 8 h after cold stress, observed in experiment [20,29,74–76].

The main feature of the compact model is kinase modularization. The kinase module makes the detailed chain of kinase cascade reaction simpler. The model allows us not to care about a network of kinases, but their regulatory relationships and functions. Biochemical events primarily focus on the positive and negative transcription factors of CBF3 or COR15A, and the negative feedback loop formed by CBF3 and ZAT12. In absence of experimental control with several genes, the predicted time evolutions (Figure 2, 5, 6) are conducive to provide advice for molecular mechanism analysis. The experimental data obtained from different plants are precious, and they render the model be practical. The prediction and data fitting of the expression of cold-responsive *CBF* and *COR* genes in different plants also further verify the effectiveness of the model. There are two experimental results worth noting. Firstly, the core gene that increases cold acclimation in different plants is not CBF3 invariably, proved to be CBF1 in tomato [75,77]. Secondly, differences in gene regulatory network are likewise verified experimentally, e.g., MYB15 is a positive transcription factor of CBF1 in tomato with the effect of light [44], and the regulatory relationship is the same as that of *Arabidopsis* in non-heading Chinese cabbage, a cruciferous plant [78].

In the core chain of sequential reactions, kinases have both positive promotion and negative inhibition, may considered as neutralized regulation, on the synthesis of phosphorylated ICE1 (Figure 1). The activated CaM would exert antagonistic effects on phosphorylated ICE1, whose degradation promoted by kinases. The subsequent biochemical events are the core part of the regulatory network, i.e., the temporal evolution of *CBF3* mRNA is activated by ICE1 and inhibited by MYB15 and ZAT12, which was induced by CBF3. Although qualitative information about the identification and mutual regulation of genes related to cold response are available, few studies incorporate these regulatory relationships into predictive models. Such models are invaluable because mathematical models and computer simulations can help us understand the inherent nature and dynamics of these processes, and make well-founded predictions about their future development and the impact of their interaction with the environment. Mathematical analysis is an effective to obtain comprehensive and global view of cold response pathway.

Cold stress medium of constructing ICE1-CBF-COR pathway is Ca^2+^, known as a second messenger response to cold shock. A prevailing kinetic model for Ca^2+^ oscillations based on the Ca^2+^-induced Ca^2+^ release mechanism is applied again in our model [60,61]. The concentration of Ca^2+^ in the cytoplasm instantly raises after cold treatment, but the different dynamics between one-compartment and two-compartment model are uncertain. In order to explore the factors that cause changes in the expression pattern of *CBF3* and *COR15A* mRNA, we simulate the time evolutions under Ca^2+^ two-compartment model in small amounts and great deal. Numerical simulations show that the amplitude of *CBF3* and *COR15A* mRNA increases with the rising number of Ca^2+^ spikes (Figure S2 and S10). Meanwhile, two-compartment Ca^2+^ model can both exert a single spike and multi-spike oscillations, which depends on the parameter *a a*nd *β (*Figure S3).

Gene mutation and overexpression are generally used to evaluate the degree of influence of wild-type genes on the expression of neighborhood genes in the transcriptome, pathway or network [79]. The model predicts the effects of mutation or overexpression of genes in the cold response pathway on *CBF3* transcription. The *ice1* mutation suppresses the transcription of CBF3 (Figure 4A). Chinnusamy et al. [52] have illustrated that the *ice1* mutation also blocks the expression of downstream genes of CBFs, while the overexpressed ICE1 in WT is an accelerant of CBF3 in transgenic plants, leading to a significant reduction and accession in plant chilling and freezing tolerance, respectively. The loss-of-function of MYB15 results in the increased *CBF3* mRNA level in the cold (Figure 4B). The *myb15* mutant plants have shown increased tolerance to low temperature [41]. Overexpression of CBF-inhibitor ZAT12 similarly hinders the expression of CBF3 (Figure 4C). On account of no light effect considered in cold response pathway, some experimental results are not reflected in the model simulation, e.g., CBF3 transcription shows a peak at 24 h (corresponding to 26 h in Figure 4A, 4C), observed in Figure 1C and Figure 10A from the study of Chinnusamy et al. [52] and Vogel et al. [43], respectively.

Desensitization avoids the plant cells suffering deeper damage from the uncontrolled stimulation, whereas resensitization allows cells to recover or maintain their responsiveness [80]. Desensitization and resensitization of *CBF3* and *COR15A* transcription have occurred in the mechanism of consecutive warm/cold stimulations [65]. Rapid cooling pulses to low temperature (4°C) causes the observably decrease (Figure 5A), slowly decline (Figure 5B) and invariable amplitudes (Figure 5C) of *CBF3* transcription after two warm/cold cycles. *COR15A* transcript levels are rapidly induced, followed by maintaining a relative steady state. The cold-sensing resensitization have been verified to recur after 24-hr at room temperatures (20°C) [65]. And then exposure to low temperature (4°C) leads to a stronger transient increase of *CBF3* mRNA level and a higher steady-state of *COR15A* mRNA level compared with those under less than 24 h warm-stage (Figure 6). While the time interval of cold stimulation increases, the desensitization and resensitization phenomenon would gradually abate (Figure 5B) and eventually disappear over time (Figure 5C). The lower limit value of temperature drop induced the sharply increasing peak of *CBF3* and the slightly higher steady-state level of *COR15A* (Figure 7).

Ordinary differential equations (ODEs) allow a better assessment of the effect of a control parameter on the dynamics of the system. However, parameterization is a great challenge for us. A small variation in the value of a parameter may cause the switch from one mode of behavior to the other [81]. A part of main parameters could be +measured by experimental control, while most parameters are not able to quantitate easily. Therefore, model parameters are selected by cost function and adjusted intuitively or comparably. Scanning a wide range of parameter values is an alternative method to explore the dynamic behavior of the model. In addition, sensitivity analysis improves the prediction of the model and examines the model quantitatively response to the variation in input variables [82–84].

The model contains 10 variables (except for Ca^2+^ and CaM) and 47 parameters. Both the above-mentioned parameter scanning and sensitivity analysis are applied in our model for the cold response pathway, explained minutely in Section 3 of Supplementary Material. Few parameter values, such as Hill coefficients [85,86], have been caught quantitatively and qualitatively by experimental control. Nonetheless, the characteristics of phosphorylation or dephosphorylation rate, enzyme-reaction Michaelis-Menten constants, and activation or inhibition thresholds have been not yet yielded.

The qualitative characteristics of *CBF3* mainly comprised the amplitude, time courses, and half-peak width. The trial-and-error procedure is carried out to meet these characteristic requirements. Hence, the process of the first step is to implement the concentration peak of *CBF3* mRNA ranging from 1 to 6 h after cold stimulation [40,41,43,52,65]. In the second step, the model need to achieve 20–500 amplitude fold-increase range compared with the prior steady state at room temperature (20°C) [52,65,87], followed by the next step through fulfilling the half-peak of the width in *CBF3* expression in the range of 3–6 h. The kinetic equations of one-compact model and two-compact model are completely consistent with those in the computational model. Therefore, the traces and settings of the kinetic parameters could be found in the discussion part and the first part of the corresponding Supplementary Material, respectively [28].

Sensitivity analysis of parameters (Figure S9) is also essential for the realization of *CBF3* transcription characteristics (see Section 3 in Supplementary Material). The model allows us to find the main parameter set affecting the characteristics of cold-elicited *CBF3* transcription, preparing for further experimental test and interpretation of biological significance.

An unobserved phenomenon in the experiment, the oscillation of *CBF3*, could occur for some choice of parameters in the model. *ZAT12* oscillated along with *CBF3*, whereas the oscillation of other genes in the network do not appear. On the one hand, this oscillation appearance might be formed by the negative feedback of CBF3-ZAT12. On the other hand, unconsidered environmental factors, such as transient cold shock, are involved in regulating cold-related genes oscillation. On top of that, circadian clock genes, such as CCA1/LHY [87], make induction of CBF3 transcription. In the absence of light, the expression of circadian clock genes is low, but the induced periodic oscillation mechanism would still exist. Photoperiod is a significant environmental factor for plant growth and development [88]. The cold response pathway model with light effect may become more complicated, which is the topic of the future research.

## Models and methods

We aim to develop a compact model for plant response to cold stress, with all kinases regarded as a kinase module, i.e., biochemical events begin from the transient Ca^2+^ increase after cold stress to the activation of CaM, followed by the increasing expression of CBF3. CBF3-regulated cold-responsive gene *COR15A* has a rising expression in the following reaction step to increase plant cold tolerance.

### Modeling influx of Ca^2+^ and activation of Calmodulin under cold shock

Based on two forms of Ca^2+^ model (one-compartment and two-compartment model), which rising in a single pulse and a cascade of spikes, the Ca^2+^ kinetic equations were derived from the previous research, i.e., a transient increase in cytosolic Ca^2+^ followed by an exponential reduction [69] and signal-induced Ca^2+^ oscillations [60,61] (See eq.(1)-(2) in Supplementary Materials). The one-compartment model is simple as it consists of only two variables. A single peak of Ca^2+^ with proper magnitude and duration are generated by selecting appropriate parameters. Goldbeter et al. [61] have discovered the sustained oscillation of cytosolic Ca^2+^ caused by the increase of inositol 1,4,5-triphosphate (InsP3) triggered by external stimuli. Releasing a certain amount of Ca^2+^ from intracellular storage occurs, followed by a subsequent increasing of cytosolic Ca^2+^.

CaM is the most fully characterized Ca^2+^ sensor in plants. Individual CaM has different spatial, temporal expression patterns, as in magnitude in response to the stimuli [89]. We directly use the activation ratio of CaM to regulate cold transduction signal.

### Model equations

Compared to the existing computational model [28] including 17 ODEs with 17 variables and 77 parameters, the compact model incorporates 13 ODEs (See Supplementary Materials) with 13 variables and 47 parameters, eq. (1)–(3) denote kinetics of Ca^2+^ one-compartment model, Ca^2+^ two-compartment model and activated CaM by Ca^2+^. The other 10 variables involving the concentration of MYB15 protein, unphosphorylated and phosphorylated ICE1 protein, *CBF3* mRNA, unphosphorylated and phosphorylated CBF3 protein, *ZAT12* mRNA, ZAT12 protein, *COR15A* mRNA and COR15A protein are listed from eq. (4) to eq. (13). All rates of protein synthesis are linear except for constant synthesis rate of ICE1 protein. All degradation rates of protein and mRNA are non-linear in terms of Michaelis-Menten kinetics. The activations/inhibitions of mRNA transcription are modeled using increment/decrement Hill-type terms, respectively. Parameter values are obtained through cost function as described in the Supplementary Materials.

## Acknowledgements

This work was financially supported by the National Natural Science Foundation of China (11871268, 11171155), National Natural Science Foundation of Jiangsu (BK20171370), the Key Projects of the National Key Research and Development Plan (2017YFD0101803), the State Key Program of the Natural Science Foundation of China (31330067), and the China Agriculture Research System (CARS-23-A-06).

## Author Contributions

**Conceptualization:**Xiong You, Ting Huang

**Funding acquisition:** Xilin Hou, Xiong You

**Investigation:** Ting Huang, Xiong You

**Methodology:** Yue Wu, Hengmin Lv

**Software:** Ting Huang, Yue Wu, Hengmin Lv, Yiting Shu, Linxuan Yu, Haoyu Yang

**Supervision:** Xilin Hou, Xiong You **Visualization:** Ting Huang, Yue Wu

**Writing – original draft:** Ting Huang

**Writing – review & editing:** Ting Huang, Yue Wu, Hengmin Lv, Yiting Shu, Linxuan Yu, Haoyu Yang, Xilin Hou, Xiong You

## Supporting information captions

**SI Text. Supplementary notes on the modeling the Ca**^**2+**^ **signal triggered by the cold stress, kinetic equations for the cold stress response pathway, model parameterization and sensitivity analysis, supplementary Figures and Tables, as well as the methods of data extraction and numerical simulation**.

